# Feasibility of phosphoproteomics on leftover samples after RNA extraction with guanidinium thiocyanate

**DOI:** 10.1101/2020.08.27.269894

**Authors:** Frank Rolfs, Sander R. Piersma, Mariana Paes Dias, Jos Jonkers, Connie R. Jimenez

## Abstract

In daily practice, different types of biomolecules are usually extracted for large-scale ‘omics’ analysis with tailored protocols. However, when sample material is limited, an all-in-one strategy is preferable. While lysis of cells and tissues with urea is the accepted standard for phosphoproteomic applications, DNA, RNA and proteins can be simultaneously extracted from small samples using acid guanidinium thiocyanate-phenol-chloroform (AGPC). Use of AGPC for mass spectrometry (MS)-based phosphoproteomics has been reported, but not benchmarked. Here we compared urea-with AGPC-based protein extraction, profiling phosphorylations in the DNA damage response pathway after ionizing irradiation of U2OS cells as proof of principle. On average we identified circa 9000 phosphosites per sample with both extraction methods. Moreover, we observed high similarity of phosphosite characteristics (e.g. 94% shared class 1 identifications) and deduced kinase activities (e.g. ATM, ATR, CHEK1/2, PRKDC). AGPC-based sample extraction can thus replace standard cell lysates for phosphoproteomic workflows and may thus be an attractive way to obtain input material for multiple omics workflows, yielding several data types from a single sample.

## Introduction

Mass spectrometry (MS)-based phosphoproteomics is a powerful tool to study cell signaling in a global fashion and with high resolution, either at steady state or following perturbation or development of disease (1). Moreover, the study of phosphorylation sites allows the identification of upstream effectors, i.e. kinases, and thus predictions on their activity (2, 3).

A commonly used and accepted method to isolate proteins from biospecimens for phosphoproteomic profiling is the application of chaotropic agents such as urea (4-7). In daily practice, other biomolecules such as nucleic acids are usually extracted using distinct protocols. However, in situations where available sample material is limited it is desirable to extract as much of the various types of biomolecules as possible using one protocol. This can be achieved through acid guanidinium thiocyanate-phenol-chloroform (AGPC) extraction using commercially available reagents like TRIzol, TriFast, TRI Reagent, or RNA-Bee (8-10). It has been shown that proteins isolated in such a way are compatible with quantitative mass spectrometry for protein expression profiling (11-13). Although AGPC-based protein extraction has been used for the analysis of the phosphoproteome (14-16), a direct comparison of this isolation method with the standard urea approach for the study of cell signaling using phosphoproteomics is currently lacking.

In this work, we compared the phosphoproteome of protein extracts obtained with a standard urea protocol or an AGPC reagent (RNA-Bee). In a functional context, we also compared coverage of the well-studied phosphorylation response after DNA damage (17), performing differential analysis of phosphorylation site data from untreated versus irradiated cells. Using high-resolution tandem mass spectrometry and recent bioinformatic tools such as inferred kinase activity (INKA) analysis and phosphosite signature enrichment analysis (PTM-SEA) (3, 18), we show that phosphorylation data for both extraction methods are highly similar.

## Experimental Procedures

### U2OS cell culture, irradiation and protein isolation

U2OS cells (KCLB Cat# 30096, RRID:CVCL_0042) were cultured in 15 cm dishes in DMEM supplemented with GlutaMAX (Gibco). Medium was supplemented with 10% FBS (Serana) and 50 units/ml penicillin-streptomycin (Gibco), and cells were grown at 37°C under normal oxygen conditions to 80-90% confluency before treatment.

Ionizing irradiation was performed using a Gammacell 40 Extractor low dose rate research irradiator (Best Theratronics Ltd.) set to a dose of 10 Gy and cells were allowed to recover for one hour before harvesting.

For phosphoprotein extraction with RNA-Bee, six cell culture dishes (3 untreated, 3 irradiated) were washed with PBS, lysed with 2 ml RNA-Bee (Tel-Test), frozen on dry ice and stored at -80°C until further use. Subsequently, a 1 ml aliquot was thawed at room temperature and 150 µl chloroform was added, the tube was shaken vigorously by hand and incubated for 5 min at room temperature. Afterwards, the sample was centrifuged (12000 x g, 15 min, 4°C). After removal of the aqeous phase containing RNA and addition of 300 µl 100% ethanol, the sample was incubated for 3 min at room temperature and centrifuged (2000 x g, 5 min, 4°C) to pellet the DNA. The resulting supernatant was precipitated with 3 ml ice-cold acetone, incubated for 5 min at room temperature and centrifuged (2800 x g, 5 min 4°C) to collect precipitated proteins. The protein pellet was washed twice with 500 µl 95% ethanol and subsequently dissolved in 800 µl urea lysis buffer (9 M urea, 20 mM HEPES pH 8.0, 1 mM sodium orthovanadate, 2.5 mM sodium pyrophosphate, 1 mM beta-glycerophosphate) by pipetting. Afterwards, samples were intensely vortexed at room temperature using an Eppendorf thermomixer for about 1-2 h. Finally, samples were sonicated three cycles (20 s on, 20 s off at maximum amplitude) using a Branson high-intensity cuphorn sonicator and cleared by centrifugation (16000 x g, 10 min, room temperature).

For phosphoprotein extraction with the standard urea protocol, another 6 cell culture dishes (3 untreated, 3 irradiated) were washed with PBS and lysed using 2 ml urea lysis buffer followed by three cycles of sonication (20 s on, 20 s off at maximum amplitude). Samples were then cleared by centrifugation (16000 x g, 10 min, room temperature). Protein concentration was determined with a Pierce BCA Protein Assay Kit (Thermo Scientific) and all samples were stored at -80°C until further use.

### Protein digestion and phosphopeptide enrichment

Experimental steps were performed as previously described (3, 19). Briefly, an equivalent of 500 µg total protein was used and diluted with urea lysis buffer to a final concentration of 1 µg/µl in a 500 µl total volume. Dithiothreitol (DTT) was added to a final concentration of 4 mM and samples were incubated for 30 min at 55°C in a water bath. After cooling down to room temperature, iodoacetamide was added to a final concentration of 10 mM and samples were incubated for 15 min in the dark. The samples were then diluted to 2 M urea final concentration via the addition of 20 mM HEPES pH 8.0 and digested overnight with 5 µg/ml sequence-modified trypsin (Promega) at room temperature.

The digests were then acidified with trifluoroacetic acid (TFA) to a final concentration of 0.1% and desalted using Oasis HLB columns (10 mg capacity, Waters). Columns were activated with acetonitrile (ACN) and equilibrated in 0.1% TFA. Bound peptides were washed twice with 0.1% TFA and eluted in 0.1 % TFA, 80% ACN solution.

Phosphopeptide enrichment of desalted peptides was performed hereafter as previously descibed (19) using TiO_2_ beads: For this, desalted peptides were diluted 1:1 with lactic acid solution (0.3 g/ml lactic acid, 0.07% TFA, 53% ACN). For phosphopeptide capture, 2.5 mg of TiO_2_ beads (GL sciences, 10 µm) were packed in Stage-tips fitted with a 16G-needle punch of C8 material (3M Empore) at the narrow end. Tips containing the TiO_2_ bed were first washed with 200 µl of 0.1% TFA/80% ACN and equilibrated with 200 µl of 300 mM lactic acid solution. Desalted peptide loading of tips was performed in 5 cycles using 200 µl portions of the peptide mixture in each cycle. The TiO_2_ bed with bound phosphopeptides was washed first with 200 µl lactic acid solution and secondly with 200 µl 0.1% TFA/80% ACN. All steps were performed via centrifugation at 1500 x g for 4 min. Afterwards, phosphopeptides were eluted in two steps with 50 µl 0.5% piperidine (Thermo Fisher Scientific) and 50 µl 5% piperidine, and subsequently quenched in 100 µl 20% H_3_PO_4_. This was followed by desalting of phosphopeptides using 200-µl Stage tips fitted with a 16G-needle punch of SDB-XC material (3M Empore) at the narrow end, which was washed with 20 µl 0.1% TFA/80% ACN and equilibrated with 20 µl 0.1% TFA. Phosphopeptides were loaded and centrifuged for 3 min at 1000 x g. SDB-XC beds were then washed with 20 µl of 0.1% TFA, and desalted phosphopeptides were eluted with 20 µl of 0.1% TFA/80% ACN. Phosphopeptides were finally dried in a vacuum centrifuge and dissolved in 20 µl 0.5% TFA/4% ACN prior to LC-MS/MS.

### LC-MS/MS

Peptides were separated using an Ultimate 3000 nanoLC-MS/MS system (Thermo Fisher Scientific) equipped with a 50 cm × 75 μm ID Acclaim Pepmap (C18, 1.9 μm) column. After injection, peptides were trapped at 3 μl/min on a 10 mm × 75 μm ID Acclaim Pepmap trap at 2% buffer B (buffer A: 0.1% formic acid (Fisher Scientific), buffer B: 80% ACN, 0.1% formic acid) and separated at 300 nl/min in a 10–40% buffer B gradient in 90 min (125 min inject-to-inject) at 35°C. Eluting peptides were ionized at a potential of +2 kVa into a Q Exactive HF mass spectrometer (Thermo Fisher Scientific). Intact masses were measured from m/z 350-1400 at resolution 120.000 (at m/z 200) in the Orbitrap using an AGC target value of 3E6 charges and a maxIT of 100 ms. The top 15 for peptide signals (charge-states 2+ and higher) were submitted to MS/MS in the HCD (higher-energy collision) cell (1.4 amu isolation width, 26% normalized collision energy). MS/MS spectra were acquired at resolution 15000 (at m/z 200) in the orbitrap using an AGC target value of 1E6 charges, a maxIT of 64 ms, and an underfill ratio of 0.1%, resulting in an intensity threshold for MS/MS of 1.3E5. Dynamic exclusion was applied with a repeat count of 1 and an exclusion time of 30 s.

### Peptide identification

MS/MS spectra were searched against the Swissprot *Homo sapiens* reference proteome (downloaded February 2019, canonical and isoforms, 42,417 entries) using MaxQuant 1.6.4.0 (20, 21) software. Enzyme specificity was set to trypsin and up to two missed cleavages were allowed. Cysteine carboxamidomethylation (Cys, +57.021464 Da) was treated as fixed modification and serine, threonine and tyrosine phosphorylation (+79.966330 Da), methionine oxidation (Met,+15.994915 Da) and N-terminal acetylation (N-terminal, +42.010565 Da) as variable modifications. Peptide precursor and fragment ions were searched with a maximum mass deviation of 4.5 ppm and 20 ppm, respectively. Peptide, protein and site identifications were filtered at an FDR of 1% using the decoy database strategy. The minimal peptide length was 7 amino acids, the minimum Andromeda score for modified peptides was 40, and the corresponding minimum delta score was 6 (default MaxQuant settings). Peptide identifications were propagated across samples with the match between runs option checked.

### Label-free phosphopeptide quantification & data analysis

Phosphopeptides were quantified by their extracted ion intensities (‘Intensity’ in MaxQuant). For phosphosites, MaxQuant output data (Phospho (STY)Sites.txt) was loaded into R (version 3.6.3) (22) and further processed using a custom script. In brief, decoy database hits, contaminants and all-zero intensity rows were excluded. Numbers of phosphosites and missing values (Supplementary Table 1 B-C) were determined by samplewise summation of data in intensity columns per phosphosite entry in the resulting table (Supplementary Table 1 A). For each site, this involves three separate intensity columns, __1/ __2/ __3, harboring data derived from phosphorylated peptides with 1, 2 or ≥ 3 phosphorylation sites, respectively, which are distinguished as they yield different signal magnitudes. The data matrix was then transformed from a wide format with separate 1, 2, and 3 columns to a long format with separate rows for PS quantifications derived from phosphopeptides with 1 (__1), 2 (__2) or ≥ 3 (__3) phosphorylation sites. Data were log2-transformed and normalized on the median intensity of all identified phosphosites (Supplementary Table 1 D). Only phosphosites with a localization probability ≥0.75 (class 1) were used for further analysis.

Correlation analysis was based on Pearson correlation of class 1 phosphosites and plotted using the ComplexHeatmap R package (23). For dimension reduction analysis, the uniform manifold approximation and projection (UMAP) algorithm (24, 25) was used as implemented in the umap R package. Calculations were based on class 1 phosphosite intensities and missing values were replaced with zeros.

Gene ontology analysis was performed using the ClueGO plug-in version 2.5.4 (26) for Cytoscape software (27) with default settings. GO-term fusion was allowed and only significant terms after p-value correction with Bonferroni step down (< 0.05) were considered.

INKA analysis was performed as described previously (3), using the online version accessible at https://inkascore.org. Downloaded INKA scores were filtered for full data presence for replicates in one of the treatment groups (either untreated or irradiated). The limma (28) R package was used to perform differential expression analysis for INKA score and phosphosite intensity data.

Furthermore, phosphosite intensity data were filtered for the presence of at least 2 out of 3 data points for the samples in at least one of the treatment groups. In case of data presence in one group and absence in the other (phosphosite on/off behaviour), only observations without missing values in the ‘phosphosite on’ group were allowed. Solely in these cases missing values were imputed in the ‘phosphosite off’ group with a zero to derive a p-value. Fold changes were determined using the mean of each treatment group and the antilog value was calculated (2^(mean of log2 values group 2 - mean of log2 values group 1)^). To rank phosphosites for PTM-SEA, the negative log10 p-values derived from limma were multiplied with the sign of the fold change. In case of duplicated phosphosite amino acid windows, the most significant (lowest p-value) entry was used. PTM-SEA was performed using the GenePattern platform (29) and output was further processed in R.

Standard deviation to estimate resemblance of phosphosites from each extraction method was calculated per treatment group from urea and RNAB derived phosphosite intensities together and missing values were imputed with zero.

### Experimental Design and Statistical Rationale

Single-shot phosphoproteomic profiling was performed for 12 samples (15 cm cell culture dishes) of the U2OS cell line. For each variation (‘urea’, extraction with urea lysis buffer; ‘RNAB’, extraction with RNA-Bee; ‘IR’, irradiation; ‘NT’, no treatment) 3 replicate samples were used. The selection of a single cell line as proof of principle minimizes inter-cell line variation and thus allows a more unbiased focus on the phosphorylation sites in each condition. Phosphoproteomic workflow reproducibility was assessed previously (19) and thus motivated our choice of three replicates per condition as a sample number which can be easily handled at a given time without compromising the quality of processing. To avoid batch effects, measurement of samples and conditions was alternated (3 cycles with one sample each of the urea-NT, urea-IR, RNAB-NT, and RNAB-IR groups). For the analysis of differential phosphosites, limma was selected as it was designed for handling of differential expression studies and smaller sample sizes (28). Statistical aspects of INKA scoring and PTM-SEA were described in the original publications (3, 18).

## Results

To compare the two extraction protocols, U2OS cells were either not treated or irradiated with 10 Gy followed by one hour recovery, before protein isolation with either RNA-Bee (RNAB) or urea (three replicates for each treatment-isolation combination, Fig. 1 A and experimental procedures). For both isolation methods, a standard TiO_2_ phosphoproteomic workflow (19) was applied (experimental procedures). In total, 13,519 phosphosites (PS) were identified with an average of 9034 PS per sample. No obvious differences in missing value numbers were found between samples extracted with urea versus RNAB (Supplementary Fig. 1 A-B and Supplementary Table 1 B-C). Identified PS highly overlapped for both extraction methods, with 94.2% shared identifications for high-confidence class 1 PS (Fig. 1 B and Supplementary Fig. 2 A). Class 1 PS proportions were found to be equal for the two different extraction methods (Fig. 1 B). Non-shared class 1 PS showed lower localization probability than shared class 1 PS (Supplementary Fig. 2 B). Furthermore, when gene names of non-shared class 1 PS were used for gene ontology analysis, no significant terms were found for extraction with RNAB, and only few terms with low gene coverage were observed for urea extraction (Supplementary Fig. 2 C). Correlation analysis of normalized data (Fig. 1 C) was able to distinguish both extraction techniques but overall showed very high similarity with a minimum correlation coefficient of 0.91 (Fig. 1 C). When the UMAP dimension reduction algorithm (24, 25) was used on these phosphosite data, the major separation occurred between non-treated versus irradiated samples, not between isolation methods (Supplementary Fig. 2 D). In summary, phosphoproteomic analyses based on AGPC-extracted protein samples fully recapitulate results obtained with protein extracts prepared via standard urea buffer lysis.

**Figure 1:**
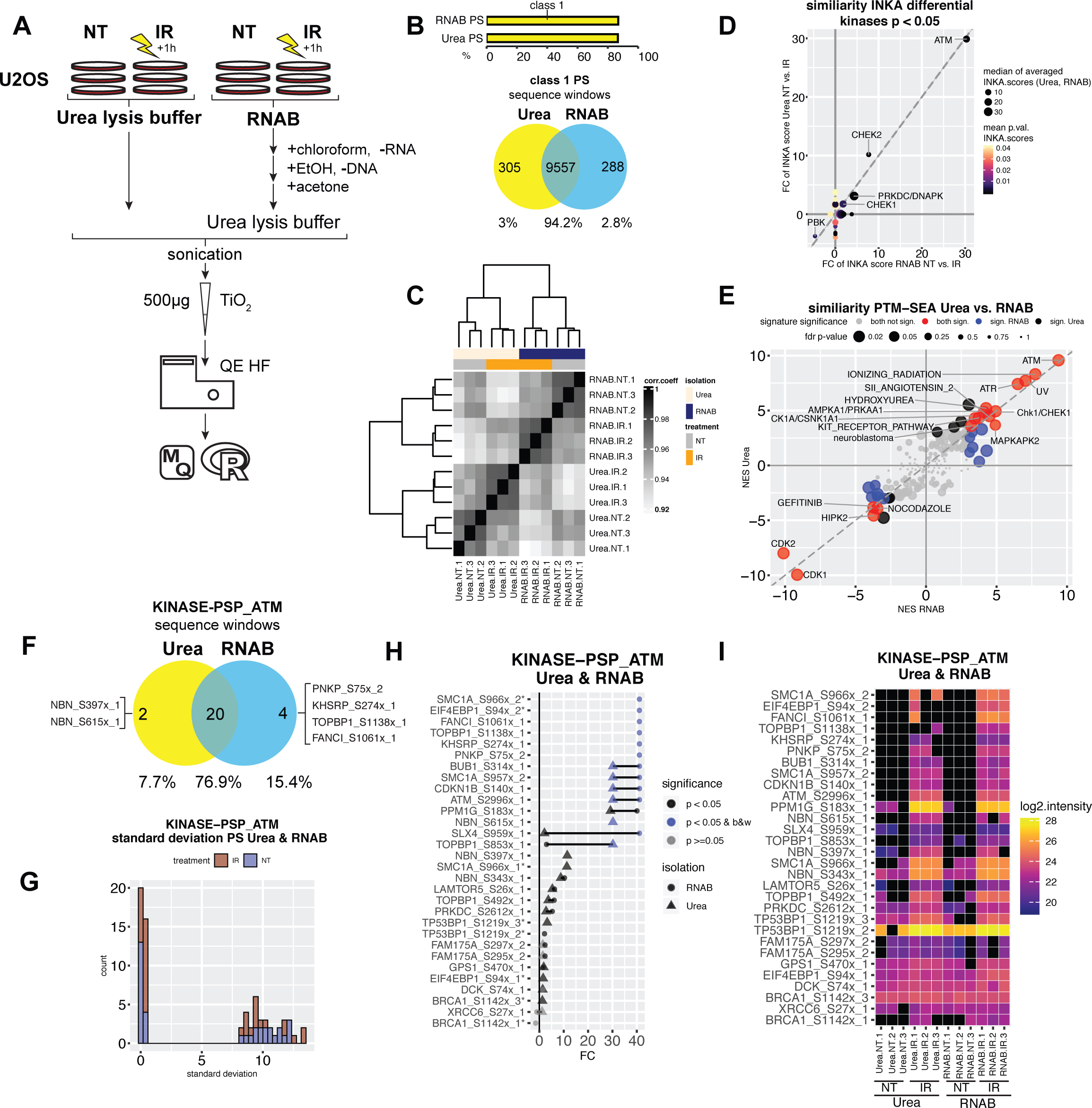
Phosphoproteomic data comparison of untreated and irradiated U2OS cells extracted with urea lysis buffer or RNA-Bee shows very high similarity. **A)** Schematic overview of the experimental setup. Three replicate 15 cm dishes with U2OS cells were irradiated with 10 Gy followed by incubation for 1 hour (IR) or left untreated (NT) and subsequently harvested with urea lysis buffer (urea) or RNA-Bee (RNAB). RNA and DNA was removed for RNAB lysed cells and proteins were precipitated with acetone and dissolved in urea lysis buffer. Lysates were sonicated and phosphopeptides enriched using TiO_2_ before measurement on a QE HF mass spectrometer. Data analysis was performed using MaxQuant and R software. **B)** Upper bar plot showing the proportion of confidently localized (class 1) phosphosites (PS) for urea and RNAB isolated samples. Lower Venn diagram showing large overlap of detected class 1 PS between urea and RNAB isolated samples. Sequence windows of identified PS were used for the overlap. **C)** Heatmap showing the correlation and strong resemblance between urea and RNAB isolated samples. Note the lowest correlation of 0.91. Pearson correlation coefficient (corr.coeff.) is based on normalized intensity data and class 1 phosphosites. **D)** Correlation of post-irradiation kinase activity for urea or RNAB lysis. Single sample inferred kinase activity (INKA) analysis was performed to detect activated kinases. Scatter plot showing INKA score fold changes (FC) of untreated (NT) vs. irradiated (IR) cells for urea or RNAB isolated specimens. Only significantly differential kinases are depicted. Limma derived p-values are color coded. INKA score magnitude is indicated via circle size and shows the median of the replicate-averaged INKA scores for urea or RNAB isolated cases for each kinase. The related individual plots are shown in Supplementary Fig. 2 E-F. **E)** Phosphosite specific signature analysis (PTM-SEA) using phosphoproteomic data of cells lysed with urea or RNAB yields highly correlated results. As a ranking metric, -log10(p-value) multiplied by the sign of the fold change was used. Scatter plot showing normalized enrichment scores (NES) of phosphosite signatures derived from untreated (NT) vs. irradiated (IR) cells after urea or RNAB isolation. Signature significance is color coded. Magnitude of p-values is indicated via circle size. P-values were only averaged for signatures that were either both significant or both non-significant with both extraction methods. Related plots showing all significant signatures separately are shown in Supplementary Fig. 2 G-H. **F)** Venn diagram showing overlap of PS found for PTM-SEA signature KINASE-PSP_ATM shown in E for urea and RNAB lysed cells. Overlap is based on sequence windows. Note that due to data presence filtering before statistical evaluation not all PS overlap here. Some of these non-shared PS were detected in a fraction of the samples, compare to H, I. Several PS are only present after irradiation (off/on behaviour); in view of the sample number, only cases with complete data presence for either irradiation or control were considered; compare to H, I. Numeric tags x_1 and x_2 indicate if PS x was derived from a singly or doubly phosphorylated peptide, respectively. **G)** Using PS overlapping with PTM-SEA signature set KINASE-PSP_ATM shown in E, the standard deviation of intensities was calculated across all samples (irrespective of isolation method) for each PS member of the signature after irradiation (IR) or no treatment (NT). The resulting frequency distribution is plotted. **H)** Cleveland dot plot showing the fold changes (FC) found for PS overlapping with the KINASE-PSP_ATM signature derived from PTM-SEA shown in E. Limma derived significance of NT vs. IR comparison is color coded. Off/on (black/white regulation) behavior of PS after irradiation is indicated. Isolation method is depicted with different shapes. Several PS are only present after irradiation, most probably due to sample number, and only cases with complete data presence for either irradiation or control were considered; compare to I. Numeric tags x_1, x_2, and x_3 indicate if PS x was derived from a singly or multiply phosphorylated peptide; equivalent PS derived from these peptides are labelled with *. **I)** Heatmap visualizing enriched PS for PTM-SEA KINASE-PSP_ATM signature shown in E. The measured log2 intensity is color coded for each PS. Black color indicates no detection.

U2OS cells display a known phosphorylation response after irradiation-induced DNA damage, and DNA damage signaling is initiated via phosphatidylinositol 3-kinase related kinases such as ATM, ATR and DNAPK/PRKDC (30). To investigate possible differences in the extent to which this response is captured with phosphoproteomic analyses of urea versus RNAB protein extracts, kinase activities were deduced for each sample using our inferred kinase activity (INKA) analysis pipeline (3). The resulting INKA scores were tested for significant differences when comparing untreated (NT) vs. irradiated (IR) samples. This analysis returned ATM, CHEK1/2 and DNAPK/PRKDC as top hits in each case (Supplementary Fig. 2 E-F and Supplementary Table 1 E), which was expected for cells one hour after irradiation (30-32). Moreover, the fold changes of these kinases showed a strong correlation between both extraction methods and were associated with similar INKA scores (Fig. 1 D).

We next used PTM-SEA (18), a phosphosite specific signature analysis tool, to identify irradiation-induced changes in signaling pathways and kinase activity in untreated vs. irradiated cells after urea or RNAB extraction. In analogy to INKA analysis, signatures for the ATM and CHEK1/2 kinases were found to be enriched after irradiation irrespective of the employed extraction method (Supplementary Fig. 2 G-H and Supplementary Table 1 F-G). Moreover, ionizing radiation-, UV- and ATR-specific signatures were enriched after irradiation, whereas cell cycle-associated CDK1/2 and nocodazole signatures were negatively enriched. Normalized enrichment scores of significant signatures for both extraction methods showed high correlation (Fig. 1 E).

Since these signatures are grouping terms, and a single PS could differ depending on the applied extraction method, we further checked phosphosites overlapping with candidate signatures of the DNA damage response as defined by the PTMsigDB repository that is used in PTM-SEA in more detail (Fig. 1 F-I and Supplementary Fig. 3). For PS of the KINASE-PSP_ATM signature, we found a strong correlation when matching enriched PS sequence windows for both extraction methods (Fig. 1 F). For each condition (NT or IR), the standard deviation jointly calculated for PS intensities of urea and RNAB extracted samples showed a peak close to zero (Fig. 1 G). In addition, the fold changes of these PS in IR vs. NT samples were in high agreement for both extraction methods (Fig. 1 H), which was also found for PS intensities (Fig 1 I). Similar observations were made for ATR, AURKB, and CDK1 signatures (KINASE-PSP_ATR, KINASE-PSP_AurB/AURKB and KINASE-PSP_CDK1, respectively; Supplementary Fig. 3 A-L).

Although the phosphorylation response to DNA damage is mainly mediated via serine/threonine protein kinases (32) and although the overall ratio of phosphotyrosine (pTyr) sites compared to serine and threonine phosphorylation sites is low (1:1800 and 1:200, respectively (33)), we also addressed possible differences in pTyr sites after extraction with urea or RNAB (Supplementary Fig. 4). We observed a small number of pTyr sites and identified on average 76 pTyr sites (0.84% of total phosphosites) per sample (Supplementary Table 1 B-C). No clear difference was observed when pTyr site intensities (Supplementary Fig. 4 A-B) or standard deviations of urea vs. RNAB PS were compared for each treatment (Supplementary Fig. 4 C).

## Discussion

Using ‘omics’ technologies, it is now possible to comprehensively quantify and characterize nearly all biological molecules that are present in a specimen. However, this requires dedicated workflows for different types of biological molecules. With small samples, it becomes a problem to subdivide them into aliquots that harbor sufficient input material for the different workflows, e.g. transcriptomics and proteomics. In these cases, it becomes inevitable to extract different types of biological molecules from the same sample. One such strategy is to use the organic phase of acid guanidinium thiocyanate-phenol-chloroform (AGPC) extracts, that are normally discarded after retrieving the RNA-containing water phase, in order to retrieve the protein complement of specimens. Although it has been shown that this approach is feasible for both proteomics (11, 12, 34) and phosphoproteomics workflows (14-16), a direct comparison of its performance relative to a dedicated (‘standard’) extraction procedure has not been reported for phosphoproteomics. Here, we show that protein extraction from organic AGPC (RNA-Bee) fractions can substitute for standard cell lysis in urea buffer in a phosphoproteomic workflow, equally well capturing central phosphoprotein players and functional biology in a use case.

For our comparison, we used the well-studied response of cells to DNA damage (30, 31, 35, 36) as a trigger of phosphorylation changes. Results obtained with the two different extraction procedures were highly similar, with over 94% of all class-1 phosphosites being shared.

To infer kinase activity following a DNA-damaging stimulus, we applied our recently developed single sample INKA algorithm (3). The deduced kinases we identified to be active 1h after irradiation, such as ATM, ATR, CHEK1/2 and DNAPK/PRKDC, are well described key kinases of DNA damage signaling and involved in the recruitment of additional downstream DNA repair factors (37, 38). We confirmed our findings with the INKA algorithm using post-translational modification signature enrichment analysis (PTM-SEA) (18) which uses signature terms that are assembled from databases (18) such as PhosphositePlus (39). Urea and RNAB derived data both reproduced results from a previous study on the systems response of U2OS cells to DNA damage at the level of the phosphoproteome, showing increased phosphorylations attributed to ATM, ATR, DNAPK/PRKDC and CHEK1/2 upon irradiation (30).

One effect of ATM, ATR and CHEK1/2 kinase activity is the inhibition of CDKs to prevent further progression through the cell cycle after DNA damage (40). For both urea and RNAB derived phosphosite data PTM-SEA identified CDK1/2 signatures as negatively enriched after irradiation. This corresponds well with two previous phosphoproteomic studies, where targets of CDK2 showed reduced phosphorylations in G361 human melanoma cells after treatment with the radiomimetic neocarzinostatin (36), or where CDK2 sequence motifs were found to be dephosphorylated after irradiation of B-lymphocyte cells (35).

PTM-SEA provides a powerful enrichment analysis approach and largely depends on current phosphorylation signature annotations for kinases, pertubations and pathways available from databases that are not complete. For more than 95% of all reported human phosphosites kinases or biological functions are unknown (1).

Like PTM-SEA, INKA analysis also captured a decrease in CDK1/2 scores that was similar for both urea and RNAB derived data (Supplementary Table 1 E), but INKA scores were not significantly differential. On the other hand, INKA analysis indicated reduced PBK activity after irradiation for both urea and RNAB data which was not detected with PTM-SEA. PBK/TOPK (PDZ-binding kinase/T-LAK cell-originated protein kinase) is involved in control of the cell cycle and mitotic progression (41). In fibrosarcoma cells it was shown that upon doxorubicin induced DNA damage and PBK overexpression, cells bypassed the G2/M DNA damage checkpoint and entered into the next mitosis (42), which might explain the need of cells to regulate PBK activity during situations of DNA damage.

Taken together, our comparison of untreated and irradiated U2OS cells reproduces previous findings on kinase activity changes upon induction of DNA damage, and identified kinase activity changes are largely overlapping for both extraction methods. We show high similarity of phosphosites identified after urea or AGPC extraction, and extend previous findings using antibodies (43). AGPC-based extraction of proteins is thus very compatible with subsequent phosphoproteome profiling studies and likely the preferred method for simultaneous isolation of multiple biomolecules such as DNA, RNA and protein/phosphoprotein when sample material is limited. AGPC-based extraction could potentially be applied in microscaling clinical biopsy workflows and in efforts on personalized multi-omic data analysis in diseases such as cancer. In this regard, recent work demonstrated the utility of a microscaled proteogenomic workflow to isolate protein, phosphoprotein, DNA and RNA from human breast cancer core needle biopsies (7). Here, samples were serially sectioned and alternating sections collected to isolate protein and nucleic acids seperately. The results of our study indicate that AGPC might enable simultaneous extraction of protein and nucleic acids from core needle biopsies. The utility of AGPC-based protein extracts for the analysis of other post-translational modifications will need to be addressed in future studies.

## Supporting information

Raw MaxQuant output data (Phospho (STY)Sites.txt)

## Acknowledgements

We thank Dr. Richard R. de Goeij-de Haas for preparation of TiO_2_ Stage-tips, Dr. Jaco C. Knol for critical reading of the manuscript and suggestions on the text. We further thank all members of the OncoProteomics and Jonkers laboratories for discussions and support during the project. This work was supported by a fellowship from the German Research Foundation (DFG) to F.R., and by grants from the Netherlands Organisation for Scientific Research (NWO-Middelgroot project 91116017 to C.R.J.), the Dutch Cancer Society (KWF project VU2013-6423) and the Oncode Institute, which is partly financed by KWF.

## Data availability

The mass spectrometry proteomics data have been deposited to the ProteomeXchange Consortium via the PRIDE (44) partner repository with the dataset identifier PXD018562.

## Author contributions

F.R. and C.R.J conceived the study. M.P.D. performed cell culture experiments, F.R. processed the samples and S.R.P. performed the MS analysis and MQ search. F.R. analyzed the data and wrote the paper together with J.J. and C.R.J. All authors contributed to finalizing the manuscript and approved the final version.

## Abbreviations

INKA: integrative inferred kinase activity
IR: irradiated
MS: mass spectrometry
NT: untreated
AGPC: acid guanidinium thiocyanate-phenol-chloroform
PS: phosphosite(s)
PTM-SEA: post-translational modification (PTM) - signature enrichment analysis
pTyr: phosphotyrosine
RNAB: RNA-Bee
U2OS: U2 osteosarcoma

## Supplementary material related to

Supplementary Figure 1 & legend

Supplementary Figure 2 & legend

Supplementary Figure 3 & legend

Supplementary Figure 4 & legend

Supplementary Table 1 legend

Supplementary file 1 legend

**Supplementary Figure 1:**
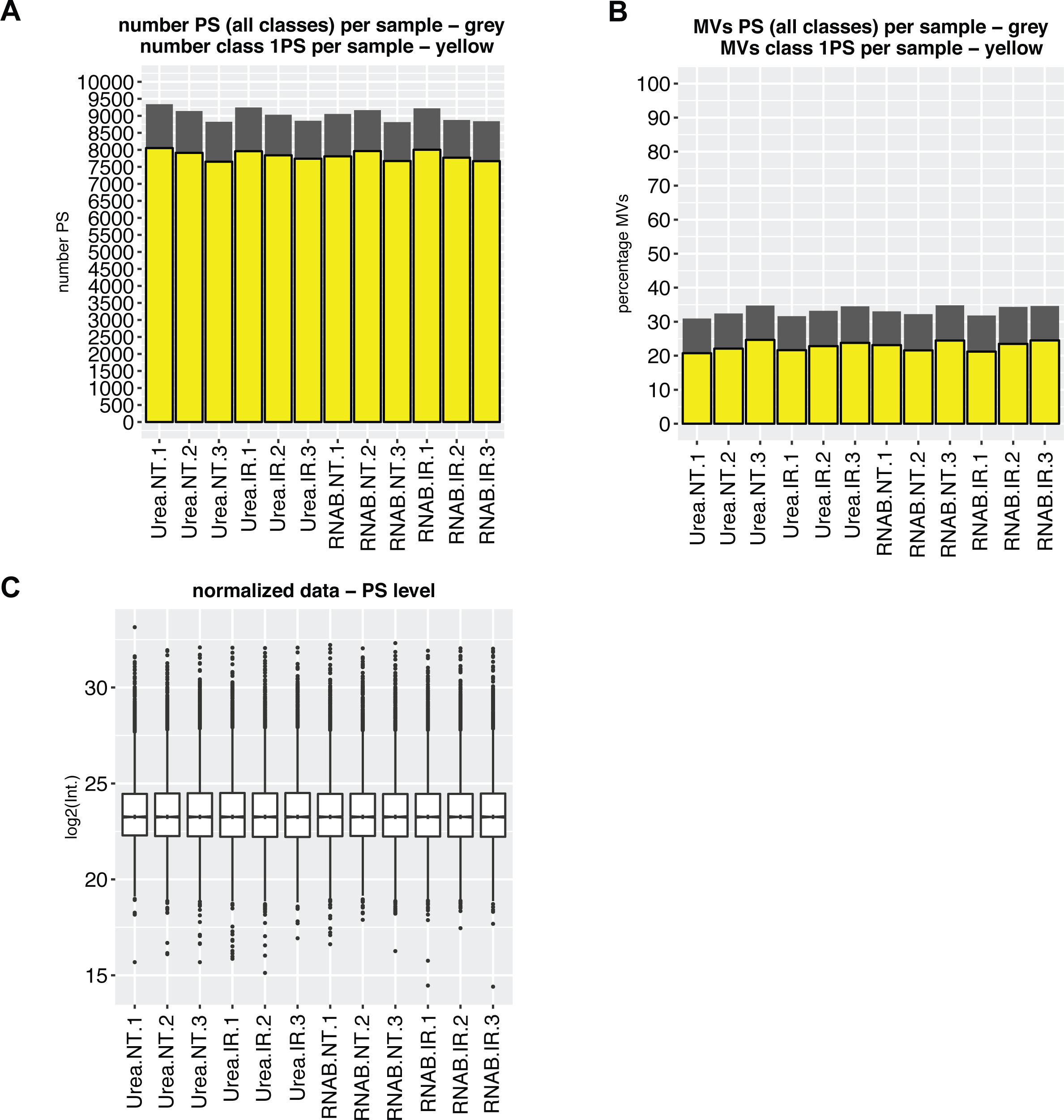
Phosphoproteomic profiling of U2OS cells after irradiation or without treatment and lysis with urea or RNA-Bee (RNAB). **A)** Bar graph depicting the number of identified phosphosites (PS) per sample. Grey color indicates all classes of identified PS. Yellow color marks the number of class 1 PS which have a localization probability ≥0.75. **B)** Bar graph showing the percentage of missing values (MV) per sample. In a data matrix of the entire experiment, MV are situations of not detected PS for a given sample which were identified in at least one of samples in the matrix. **C)** Phosphosite level box plot showing normalized data for each sample. The graph is based on PS of all classes.

**Supplementary Figure 2:**
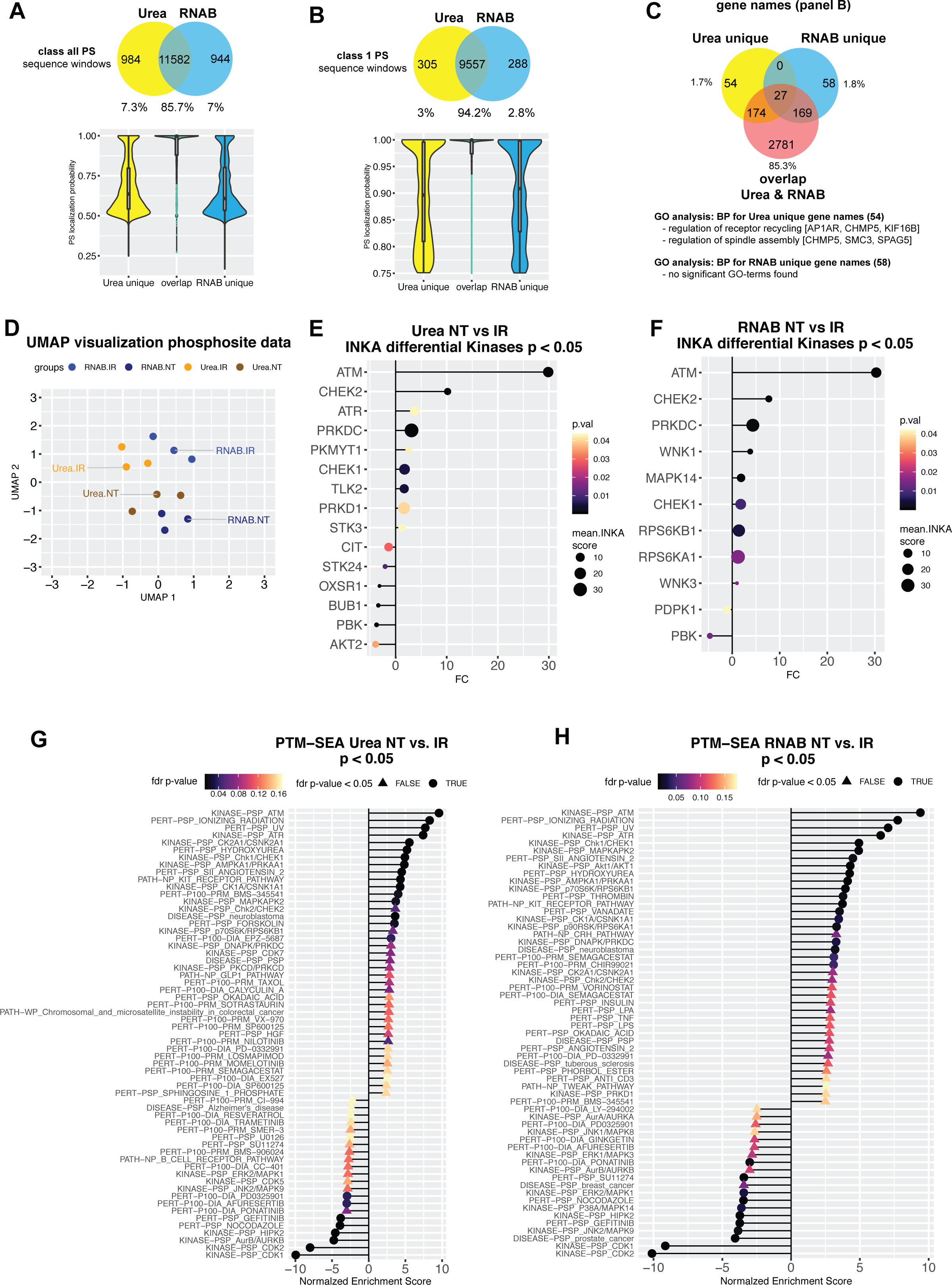
Overlay of phosphosites identified per isolation method, kinase activity and phosphosite signatures after irradiation of urea and RNA-Bee (RNAB) lysed cells. **A)** Related to Fig. 1 B. Upper Venn diagram showing overlap of class all detected phosphosites (PS) between urea and RNAB isolated samples. Sequence windows of identified PS were used. Lower violin plot illustrating localization probability distribution of PS compared in upper Venn diagram. In general, shared PS showed higher localization probability than isolation method unique PS. **B)** Related to Fig. 1 B. Same as Suppl. Fig. 2 A for class 1 phosphosites. **C)** Related to Fig. 1 B. Upper Venn diagram showing shared gene names of phosphorylated proteins shown in Suppl. Fig. 2 B. Lower part showing the result of gene ontology (GO) enrichment analysis for isolation method unique gene names using biological process (BP) terms. **D)** Related to Fig. 1 C. UMAP embedding of class 1 phosphosite data. **E)** Related to Fig. 1 D. Single sample inferred kinase activity (INKA) analysis was performed to detect activated kinases for urea lysed cells. Comparing untreated (NT) vs. irradiated (IR) samples, differential INKA scores were then detected using limma. Fold changes of significant kinases found are plotted. Significance is color coded. INKA score magnitude is indicated via circle size and shows the average INKA scores of three samples with the same treatment and isolation method. **F)** Same as E for RNAB lysed cells. **G)** Related to Fig. 1 E. Phosphosite specific signature analysis (PTM-SEA) was performed using PS of urea lysed cells and comparing untreated (NT) vs. irradiated (IR) samples. Bar plot shows normalized enrichment scores (NES) for detected, significant signatures. Adjusted p-value is color and shape coded. **H)** Same as G for RNAB lysed cells.

**Supplementary Figure 3:**
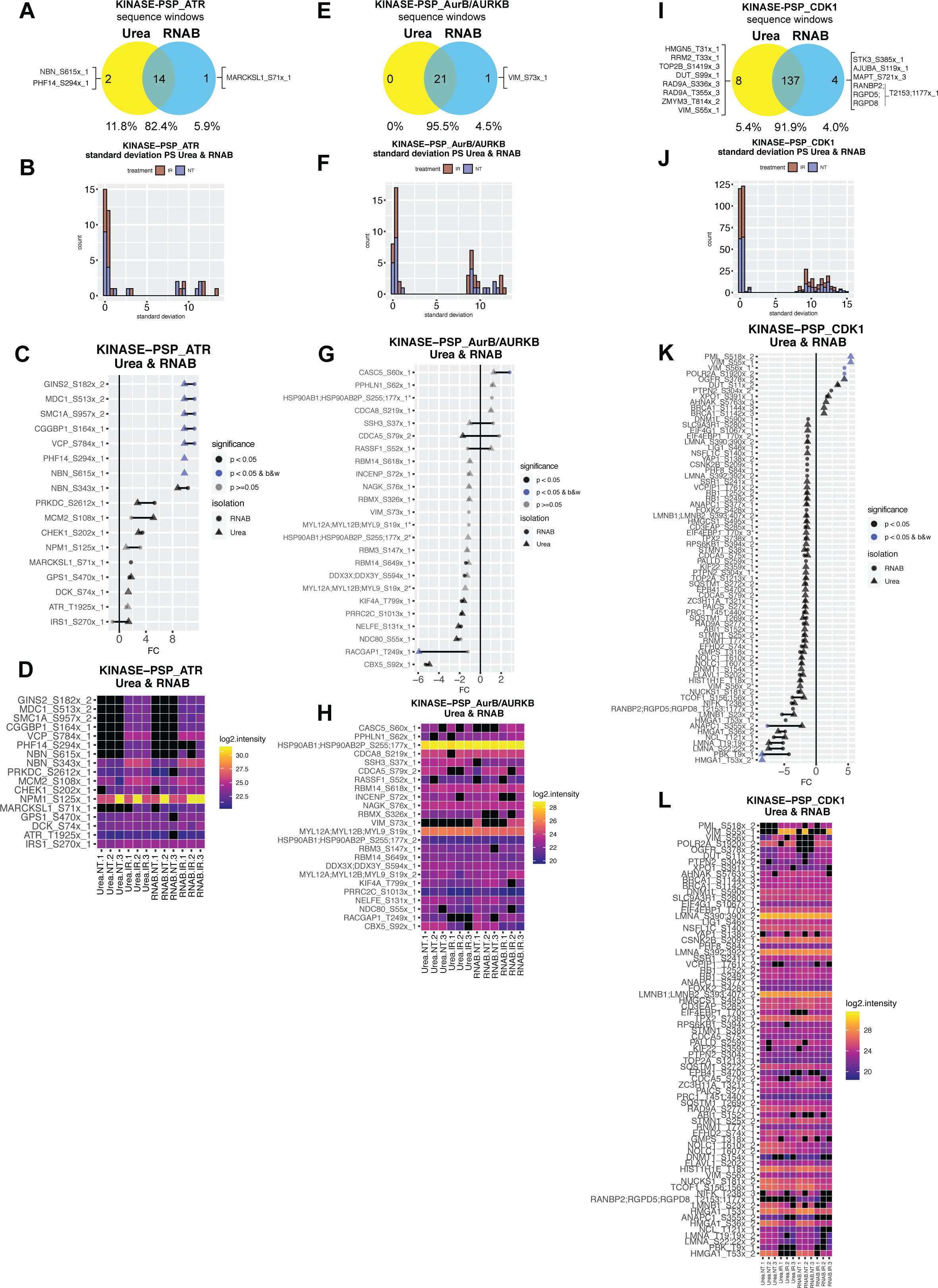
Similarity of selected PTM-SEA signatures derived from comparison of untreated vs. irradiated urea and RNA-Bee (RNAB) lysed cells. A-L) Same as Fig. 1 F-I for PTM-SEA signature KINASE-PSP_ATR (A-D), KINASE-PSP_AurB/AURKB (E-H) and KINASE-PSP_CDK1 (I-L). Note that due to the high number of signature PS for CDK1, only significant PS are depicted in K-L.

**Supplementary Figure 4:**
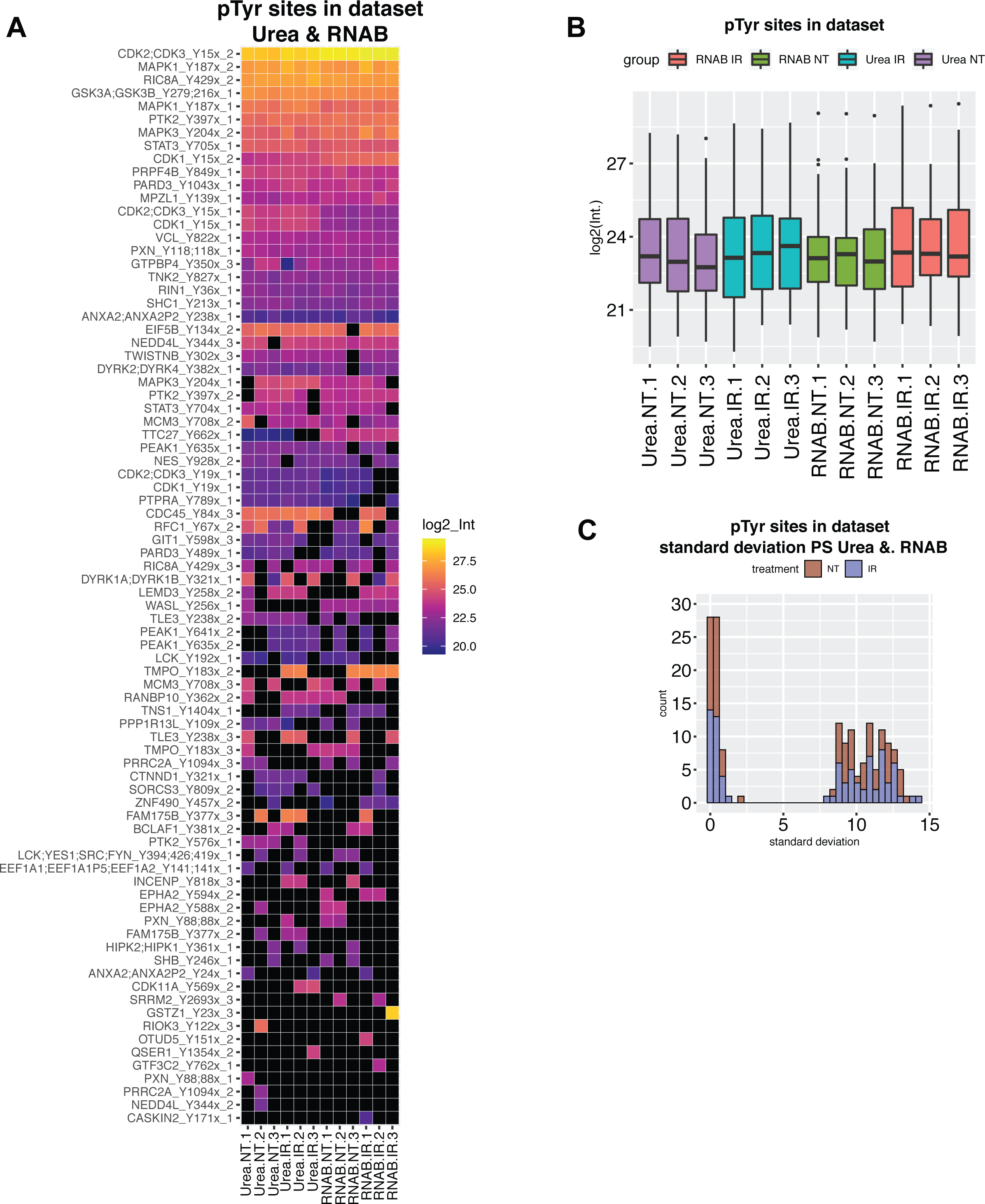
Comparison of pTyr sites detected in dataset. **A)** Heatmap visualizing class 1 pTyr sites detected in this study. The measured log2 intensity is color coded for each PS with highest intensities on top. Black color indicates no detection. x_1, x_2, x_3 indicate if PS were derived from single or multiple phosphorylated peptides. **B)** Box plot showing the combined intensities for detected class 1 pTyr sites per sample. Treatment groups and isolation methods are color coded. **C)** Standard deviation was calculated over urea and RNAB sites after irradiation (IR) or no treatment (NT). The resulting frequency distribution for class 1 pTyr sites in the dataset is plotted.

## Legend supplementary Table 1 (separate Excel file)

**A)** Shortened MaxQuant output after exclusion of reverse hits, contaminants and zero-sum rows. Basis for determination of phosphosite numbers and missing values.

**B)** Samplewise determined number of all class phosphosites, missing values and their percentages. Based on A.

**C)** Samplewise determined number of class 1 phosphosites, missing values and their percentages. Based on A.

**D)** Converted data matrix A, reshaped to account for the multiplicity of phosphorylations. Data were log2 transformed and normalized.

**E)** Scores from inferred kinase activity (INKA) analysis.

**F)** Phosphosite specific signature analysis (PTM-SEA) result for comparison of untreated and irradiated urea lysed U2OS cells.

**G)** Phosphosite specific signature analysis (PTM-SEA) result for comparison of untreated and irradiated RNAB lysed U2OS cells.

## Legend supplementary file 1 (separate text file)

Raw MaxQuant output data (Phospho (STY)Sites.txt). An overview of the samples used and their labels can be found in supplementary table 1.

